# Bridging Theory and Experiments of Priority Effects

**DOI:** 10.1101/2022.12.05.519211

**Authors:** Heng-Xing Zou, Volker H. W. Rudolf

## Abstract

Priority effects play a key role in structuring natural communities, but considerable confusion remains about how they affect different ecological systems. Synthesizing previous studies, we show that this confusion arises because the mechanisms driving priority and the temporal scale at which they operate differ among studies, leading to divergent outcomes in species interactions and biodiversity patterns. We suggest grouping priority effects into two functional categories based on their mechanisms: “frequency-dependent” priority effects that arise from positive frequency dependence, and “trait-dependent” priority effects that arise from time-dependent changes in interacting traits. Through easy quantification of these categories from experiments, we can construct community models representing diverse biological mechanisms and interactions with priority effects, therefore better predicting their consequences across ecosystems.

## Glossary

**Community assembly:** Changes in species number, identity, and abundance over time in an ecological community.

**Historical contingency:** Scenarios when the order and timing of any biotic or abiotic events affect community assembly. Priority effects can be drivers of historical contingency, but not all historical contingencies are driven by priority effects.

**Generation time:** The average time interval between an individual’s birth and the birth of its offspring in a population.

**Phenology:** Seasonal timing of life history events, e.g., breeding, migration, germination, leaf out, and recovery from hibernation.

**Positive frequency dependence:** The species with a higher frequency in the community have a higher population growth rate.

**Priority effects:** Scenarios where the outcomes of species interaction depend on their relative arrival times or initial abundances.

**Season:** A period marked by regular environmental changes that affect vital processes of species. Seasons often refer to spring, summer, fall, and winter, but they can also describe a period with suitable habitats, e.g., when ephemeral ponds have water. Communities often reassemble at the beginning of the season.

## Priority Effects: Definition, Importance, and Issues

The outcome of species interaction often depends on the temporal sequence of community **assembly** (see **Glossary**): This phenomenon, termed the “**priority effect**,” is found across a wide range of taxa and ecosystems [1–4]. By altering species interactions, priority effects can affect community composition [5–7], consequently driving biodiversity patterns [8,9] and ecosystem functioning [10–12]. Despite the long history of research into priority effects in community ecology, uncertainty remains about when and how they affect various ecological systems [7]. Much of this uncertainty arises because priority effects are used broadly to describe any relationship between the temporal sequence of arrival orders and changes in the outcomes of interactions or community states, regardless of the underlying biology and temporal scales at which the system operates. Resolving this uncertainty is not only essential to understanding the dynamics and composition of natural communities but also key to predicting how communities will change in the future as climate change reshuffles species **phenology** and the timing of species interactions over **seasons** [13–15].

Here, we review current progress in priority effects and provide a conceptual framework to understand, interpret, and predict their consequences across different systems. We first classify the diverse mechanisms causing priority effects into two categories based on the temporal scale at which they occur. We then use this framework to explain current discrepancies across empirical and theoretical studies. Finally, we provide guidelines for evaluating priority effects across different temporal and organizational scales.

## Unpacking Priority Effects: Mechanisms and Scales

Ever since the introduction of the term [16], priority effects have received much attention for their roles in shaping community assembly, such as the presence of alternative states [17,18] and **historical contingency** [19]. While the latter is often associated with priority effects, historical contingency can arise from multiple other abiotic mechanisms (e.g., dispersal, species sorting, environmental heterogeneity, disturbance [9,20–23]), many of which can also cause priority effects (see [7] for a thorough review). Here, instead of abiotic mechanisms of priority effects, we focus on the multitude of biological causes identified by experiments across many systems (Box 1; Table S1), which often differ in underlying mechanisms and temporal scales at which priority effects are measured.

### A Multitude of Mechanisms

The biological mechanisms of documented priority effects are often either unknown, indirectly inferred, or appear specific to a given study system (Table S1). For instance, in many microbial systems, priority effects are only measured by changes in community composition to differences in arrival times. While some studies point to more resolved mechanisms such as sugar concentration or pH [3,27], others are often attributed to general factors like resource preemption [24] or phylogenetic relatedness [25,26], with the specific mechanisms often unknown. In coral reef fish, priority effects often arise because early arrivers become more aggressive toward late arrivers [28–30]. Intertidal invertebrates that colonize first prevent the colonization of late-comers by physically blocking the limited substrate surface [31]. In dragonfly and damselfly communities, priority effects can be driven by intraguild predation because early arrivers gain a size advantage that allows them to prey on smaller late arrivers [32,33]. In plants, priority effects can arise from a range of different mechanisms, including competition for light [34] or pollinators [35], and plant-soil feedback (e.g., via shifts in microbiome or allelopathy) [36]. In host-pathogen systems, priority effects between two parasites co-infecting the same host can be mediated via an immune response of the host (e.g., cross-immunity) [37–39]. This multitude of mechanisms will likely lead to different consequences of priority effects in specific systems.

### A Dichotomy of Temporal Scales

Given the diversity of study systems, it is not surprising that priority effects have been studied at different temporal scales. While roughly half of the experiments we reviewed lasted more than five generations of their study organisms, the other half lasted for only one generation or less (Box 1). The two distinctive groups reveal a clear dichotomy that largely reflects differences in the life history of focal organisms. Studies that focused on organisms with relatively long **generation times**, such as plants and amphibians [1,40–42], usually last less than one generation. Therefore, the differences in arrival times are usually less than the length of their full life cycle (Figure 1) and reflect species phenology (i.e., seasonal timing within a year). Consequently, these studies largely focus on changes in traits (morphology, development or survival rates, etc.) of individuals and investigate specific biological mechanisms that cause priority effects [30,32,43]. But they do not directly measure the long-term consequences of priority effects which can only be inferred by models of population dynamics parameterized by empirical data [2,42,44–46].

**Figure 1:**
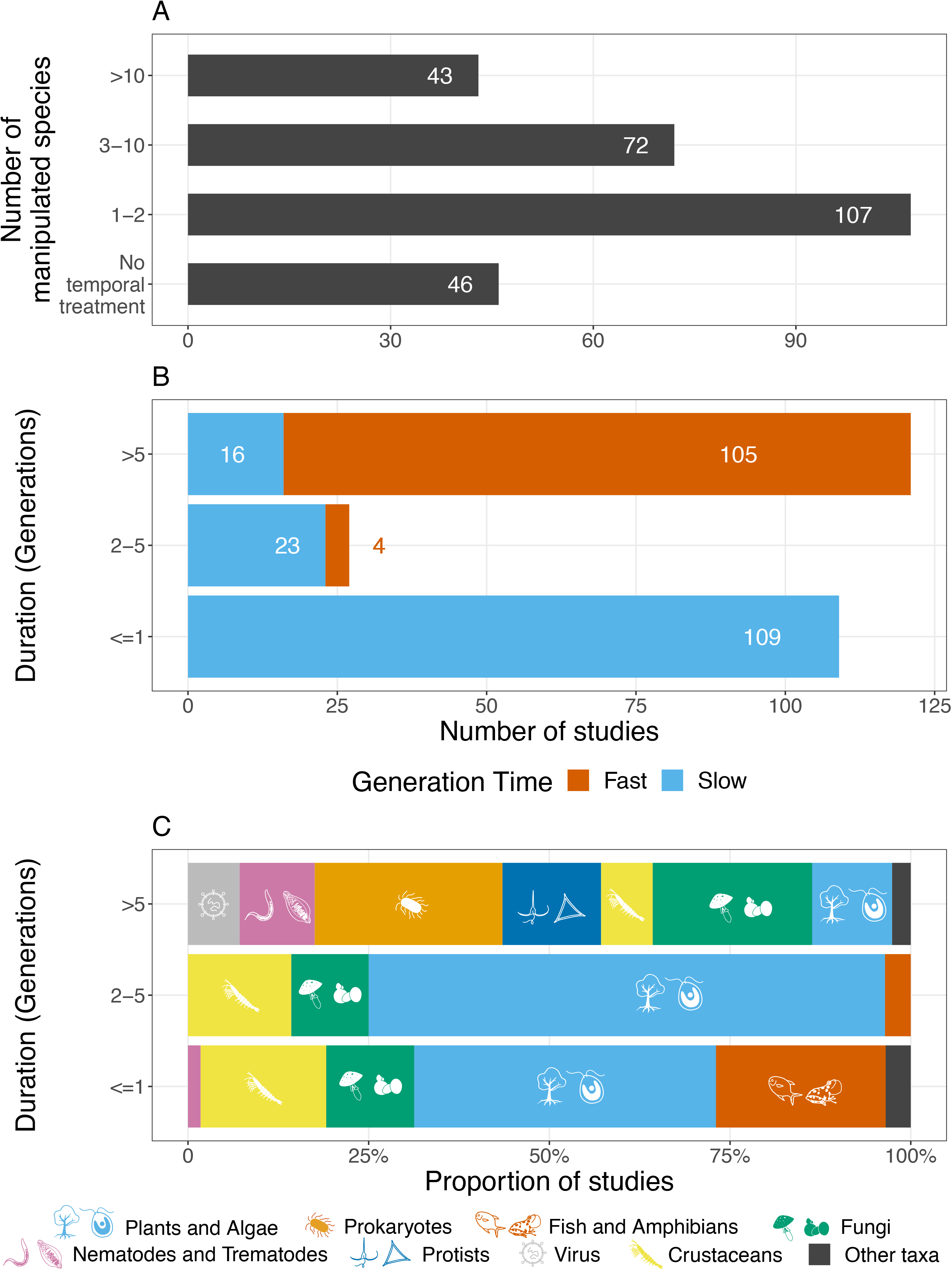
A framework of trait- and frequency-dependent priority effects. **(A)** Trait-dependent priority effects often emerge from differences in arrival times within one generation, while frequency-dependent priority effects often emerge from a numeric advantage of the early species, established by arriving multiple generations earlier. **(B)** – **(C)** With trait-dependent priority effects, the timing of species interactions affects their interaction strengths, visualized by the relationship between per-capita interspecific interaction coefficients and relative arrival times. The relationship between interactions and relative arrival times is typically either continuously nonlinear or in the extreme case piecewise (see Box 2 for examples). With frequency-dependent priority effects, per-capita interactions between species are not affected by their relative arrival times. **(D)** Trait-dependent priority effects allow for shifts between all possible outcomes of interactions, while frequency-dependent priority effects only lead to shifts between to two alternative states (e.g., which species eliminates the other, Box 2).

On the other hand, studies on organisms with short generation times such as zooplankton and microbes focus on population and community dynamics and thus often run for tens to hundreds of generations [3,47,48]. In these studies, arrival times between species typically differ by multiple generations, allowing them to study the long-term consequences of priority effects on communities (e.g., alternative stable states) [9,49,50]. However, because of the fast generation time, they typically do not investigate changes in traits of participating organisms and instead focus on environmental mechanisms that promote priority effects, such as disturbance, dispersal, resource levels, presence of predators, and pH [3,8,20,27,48].

Because of the dichotomy of temporal scales, both groups measure different response variables that are recorded at different scales (individual- vs. population-level). Therefore, patterns or mechanisms identified in studies that focus on the individual short-term (within one generation) scale may not emerge or be relevant for dynamics and patterns observed in studies on long-term community dynamics.

## The Gap Between Theoretical Framework and Empirical Studies

The theory of priority effects dates back to competition models by Lotka and Volterra, who showed that when two species in the community limit each other more than themselves, the outcome of competition is determined by the relative abundance of competitors, and the system exhibits alternative stable states [51]. Therefore, whichever species arrives earlier can gain a numeric advantage and exclude the other, leading to different final community states. This type of priority effect has also been described in the context of modern coexistence theory [52–54]. Although intuitive, this theory and its recent extensions assume that priority effects arise from a shift between alternative stable states driven by a change in the relative numeric abundances (frequencies) of species (Supplementary Material Appenidix II). This naturally raises the question: to which empirical studies does this theory apply?

We can answer this question by focusing on the key requirements for applying this mathematical framework to priority effects: (1) early arrival must provide the species with a numeric advantage when the interaction starts, (2) the combination of interaction coefficients needs to ensure **positive frequency dependence** to promote alternative stable states, and (3) the inter-(and intra-) specific per-capita interactions and demographic rates of species are fixed and independent of arrival times and sequence. The last condition effectively decouples arrival time from species interactions. It highlights that timing only matters when early arrival also ensures a numeric advantage, but the numeric advantage does not necessarily require early arrival. In fact, many studies simulate this type of priority effect without manipulating the arrival times and instead simply start experiments with an initial numeric advantage [9,55,56].

A closer inspection of empirical studies on priority effects quickly reveals that these conditions are met in some studies but not in most. Systems with short generation times such as zooplankton or microbial community are the most likely to meet these conditions because arrival times are typically separated by multiple generations, giving the early arriver enough time to reproduce, eventually outnumbering late arrivers [3,50]. However, per-capita effects are often not quantified in these studies and still could change between arrival times, e.g., due to habitat modification, phenotypic plasticity, or rapid evolution [27,57–59].

In contrast, most short-term studies documenting priority effects in empirical systems with longer generation times such as terrestrial plants, fish, and amphibians, do not satisfy these conditions. First, differences in arrival times are often too short (less than a generation) for a numeric advantage through reproduction, and many experiments keep initial densities constant across arrival time treatments. Second, studies often report that interactions in these systems are not constant but change with the timing of arrival (Table S1). Early arrival advantages caused by aggression [29,30], behavioral interference [40,60], and plant-soil feedback [36,43,61,62] are all mediated by changes in specific traits that alter the interaction between the early and the late arriver. These changes have been further quantified in insects, vertebrates, and plants [2,42,46]. These common changes in demographic rates with relative arrival time represent a mechanism of priority effects that is fundamentally different from frequency-dependent priority effects. Traditional theory based on positive frequency dependence does not explicitly include these changes in traits, and therefore cannot be applied to interpret or predict the long-term effects of the full diversity of different priority effects in nature. This gap between theory and empirical work clearly highlights current limitations and the need for a conceptual framework that captures this diversity of biological mechanisms.

## Bridging the Gap: A Unifying Framework of Priority Effects

Here, we propose a unified conceptual framework with new categorizations of priority effects to bridge the gap between theory and nature. Priority effects are defined as scenarios where the *outcomes of species interactions depend on their relative arrival times* [7]. To understand how this is possible, consider a generic two-species competition model (e.g., Beverton-Holt model; Supplementary Material Appendix II). In this system, one species can only affect the other in two ways: its density, and per-capita interaction coefficients. However, the possible equilibrium states depend only on the interaction coefficients, and only if they promote alternative stable states does the realized outcome depend on the relative frequencies of both species. Therefore, priority effects can only alter long-term outcomes when differences in arrival times also alter (1) interaction coefficients or (2) the relative frequencies of both species. Considering these facts, we suggest grouping mechanisms of priority effects into two general types: (1) “trait-dependent” priority effects which arise when differences in arrival time result in changes in per-capita interaction coefficients (Box 2), and (2) “frequency-dependent” priority effects (Table 1) which arise when differences in arrival time allow the early arriver to increase in relative frequency, leading to a priority effect that is maintained by positive frequency dependence. Note that the concept of trait-dependent priority effects includes previous concepts such as size-or stage-mediated priority effects [4,32], but is intentionally broader to include any possible trait that influences species interactions in the local community and beyond [7]. Frequency-dependent priority effects represent those studied in most traditional theories and have the same requirements (see [53] and Supplementary Material Appendix II). Note that positive frequency dependence can also arise from processes other than reproduction: In plants and slow-growing fungi, higher initial biomass often confers a higher growth rate; empirical studies on these systems often measure biomass, not fecundity, as a proxy for fitness [63,64].

Distinguishing between both types of priority effects is crucial because they have fundamentally different effects on the dynamic of the system. Trait-dependent priority effects lead to concurrent changes in key parameters that determine the possible outcome of interactions, e.g., competing species’ fitness ratio and stabilization potential (Box 2).

Depending on the original and shifted position of the interaction, this may simply mean a change in the relative frequencies of coexisting species, or a dramatic shift, e.g., from stable coexistence to competitive exclusion (Box 2). This sets them fundamentally apart from frequency-dependent priority effects, which arise from a shift between two alternative stable states (i.e., either species wins) that is only mediated by changes in their initial frequencies. Unlike frequency-dependent priority effects, trait-dependent priority effects are possible for any combination of per-capita interactions, and they can arise within a generation and therefore help explain priority effects in empirical systems with longer generation times where arrival times are separated by less than a generation.

Note that mechanisms leading to frequency-and trait-dependent priority effects can co-occur in the same system but operate at different time scales. For example, different biological mechanisms of resource competition can lead to either frequency-or trait-dependent priority effects (Box 3). In another example, the early arriving bacteria in flower nectar lowers the pH of the environment, impeding successful colonization by yeast [27]. This can be categorized as trait-dependent priority effect because arriving late changes the yeast’s demographic rates. However, on a larger temporal scale, the pH alteration can further promote bacterial growth, resulting in positive frequency dependence.

The concepts of niche preemption and niche modification (*sensu* [7]) are broad generalizations of specific mechanisms that can be categorized and modeled as frequency-or trait-dependent priority effects. In the previous example, because the relative arrival times change the demographic rates of the yeast, the priority effect arise from niche modification and is trait-dependent. Within the microbiome hosted by a single polyp of *Hydra vulgaris*, one strain of bacteria gains dominance by a higher initial abundance [65]; this is an example of niche preemption but the underlying mechanism is positive frequency-dependent. When fish first occupy a nesting site in coral reef and defend late colonizers [28], the priority effect again matches niche preemption but the driving mechanism is trait-dependent because the order of arrival changes behaviors, and the outcomes also depend on traits like body conditions [30].

Our framework could even be extended to evolutionary time scales. If one species arrives sufficiently early and adapts to local conditions, it could dominate the habitat and prevent other species or even ancestral conspecifics from colonizing, i.e., the monopolization effect [57,66–69]. At extremely large temporal scales, evolution can lead to a “phylogenetic priority effect”: early-arriving taxa could diversify and dominate in the local community [70–72].

These evolutionary priority effects can again arise from a combination of frequency-and trait-dependent mechanisms: during the adaptation process, the early colonizer can also modify the environment to lower the fitness of late colonizers (trait-dependent) or grow in large populations to achieve positive frequency dependence.

## Why It Matters

Our framework bridges the gap between theory and empirical work by providing a straightforward pathway to integrate a diversity of priority effects observed in natural systems into theoretical models (Box 2). This differentiation also allows researchers to critically evaluate the role of priority effects across a wider range of natural systems, reconciling the dichotomy between experiments on systems operating on different time scales (long- vs. short generation times; Box 1). Trait-dependent priority effects expand existing theory by including less-discussed biological mechanisms, such as evolutionary changes in species traits and fitness [57,59], and facilitative effects from early arrivers [73,74]. The latter is especially important because frequency-dependent priority effects, by definition, are only present in competitive interactions, and therefore cannot characterize the widespread positive interactions that could change with relative arrival times among species.

Furthermore, differentiating between the mechanisms of priority effects is essential to understanding the dynamics of natural systems and how they will respond to changes in the temporal structure (e.g., phenological shifts) or other types of perturbations. We already outlined how the type of mechanisms determines the consequences of shifts in arrival time for local populations, but these differences in local dynamics also scale up to drive differences in regional patterns. For instance, trait-dependent priority effects can limit the homogenizing effect of high dispersal in metacommunities because competitive hierarchies depend on species’ arrival times and not relative frequencies [8,75,76]. The mechanism of priority effects could also determine management decisions. For example, restoring a system that exhibits alternative stable states due to frequency-dependent priority effects might require manipulations of relative frequencies of species, while restoring a system driven by trait-dependent priority effects might require manipulations of actual timing [73,77]. Our categorization provides a more applicable theoretical framework of priority effects to empirical scenarios.

## Looking Forward

### Realistic Models of Priority Effects

Our classification framework emphasizes the importance of biological realism in research on priority effects, especially in theoretical studies. Models that do not capture the key mechanisms driving population and community dynamics may not accurately predict how they change over time or respond to perturbations. Thus, a realistic model of priority effects should include two key components: a base demographic model, and a function that describes how interacting species traits change with relative arrival times [15]. This integration captures the diversity of priority effects in natural systems and helps to isolate individual and synergistic effects of arrival times within and across generations.

This integration comes with new opportunities and challenges. Different mechanisms typically operate at different time scales (e.g., within vs. between generations). How models capture these time scales will ultimately depend on the life history of species, which is directly related to the temporal scales of arrival time differences (Box 1). Previous studies suggest an easy implementation in annual systems by modeling interaction coefficients as a function of arrival times (e.g., the interaction-phenology function [15]; Box 2), which allows researchers to maintain much of the original model structure and analytical tools for analyses. However, this approach does not easily extend to systems where interactions last over multiple generations within a year. These systems require different model formulations, like size/stage-structured models or agent-based models [78]. In addition, variations in arrival times are widespread in nature (e.g., between years and patches [14,79]), and individual and combined effects of different types of priority effects could lead to different community dynamics and biodiversity patterns [21,75,76]. We hope that our framework will stimulate a range of new theoretical studies that will shed new light on the different consequences of priority effects across biological systems.

### Better Quantification of Different Priority Effects

Traditionally, studies focused on the detection of priority effects by comparing dynamics that arise from different arrival orders (e.g., early vs. late treatment; Box 1). However, this simple approach is unreliable to detect the potential for priority effects. For instance, with frequency-dependent priority effects, whether the early species excludes the later arriver depends on whether its initial population is large enough or has enough time to increase relative frequency to gain a sufficient numeric advantage (Box 3). Therefore, the community dynamics depend not only on the arrival *order* but also on the arrival *time*.

To overcome these issues, future experimental research should aim to characterize priority effects across a wider range of arrival times to quantify how interactions or outcomes scale with relative arrival time (e.g., interaction-phenology function; Box 2). This would allow the experiment to uncover an accurate representation of priority effects in the focal system even when knowing the biological mechanism is difficult. The range of arrival times should scale with the life history of species to capture differences in arrival times that cover at least one generation (or length of a life stage). Relative arrival times can either be directly manipulated or simulated by competing different combinations of stages/size classes of species [2,32,42]. The effect of timing can then be measured by vital rates such as survival, growth rate, and fecundity. For systems without apparent size-or stage-mediated competition (e.g., bacteria, single-cell fungi, protists), interspecific competition coefficients can be directly evaluated from invasion growth rates [3,80]. Differences in arrival times may nevertheless allow the early arriver to establish a numeric advantage in these systems, potentially requiring crossing different initial frequencies of each species to the gradient of arrival times. The quantified interaction-phenology relationship can then be used to model long-term community dynamics under other conditions.

Although the interaction-phenology function is a powerful tool to characterize priority effects, it is phenomenological and cannot detect the potential underlying biological mechanism, such as size-mediated predation, resource competition, or environmental factors. Additional measurements on other parameters, such as functional traits, demographic parameters, or behavioral assays are required to identify the exact mechanism of priority effects in a system. Different arrival times are also likely to influence intraspecific competition in some systems with distinct ontogeny [81–83]; therefore, characterizing the relationship between intraspecific competition and arrival time might also be necessary.

### Concluding Remarks

Priority effects represent a time-explicit understanding of community dynamics, which remains one of the understudied frontiers in community ecology [15,84]. We show that by accounting for the specific mechanisms that link the relative arrival times of species to their interactions, we can extend our current theoretical framework of priority effects to cover a wider range of taxa and ecological conditions. Our framework provides new paths for modeling priority effects and highlights the need for new experimental designs that manipulate arrival times at finer temporal scales. We hope that this framework provides new perspectives on classic themes of community assembly (e.g., alternative assembly paths [85,86], repeated colonization [86,87], regional species and trait compositions [88,89], and environmental heterogeneity [90,91]), and marks the first step in considering timing in classic and burgeoning concepts in ecology (e.g., phenotypic plasticity [58,92,93], metacommunity [21,75,76], and higher-order interactions [94,95]; **Outstanding Questions**). Resolving and predicting species interactions in a temporally explicit context will be a powerful tool to decipher how natural communities respond to widespread phenological shifts under climate change.

## Acknowledgments

We thank Joshua Fowler, Tadashi Fukami, and Hannah Yin for their comments and discussions on the manuscript. We thank members of the sPriority working group funded by the Synthesis Center of the German Research Centre for Integrative Biodiversity Research (iDiv, Germany) for sharing their compiled literature of experiments on priority effects. Funding was provided by NSF DEB-1655626.

## Box 1. Empirical studies of priority effects: a literature review

We compiled 268 empirical studies of priority effects (see Supplementary Material Appendix I for methods). Despite the diverse study systems in our list of experiments, most studies were conducted on terrestrial plants (79, or 29.5%). Among all experiments, 46 (17.1%) did not explicitly manipulate arrival times but rather used initial community composition or length of natural colonization as proxies. Even among the remaining 222 studies that considered arrival time as a treatment, only 58 used a temporal gradient rather than a simple early/late/simultaneous arrival treatment (Figure IA).

We further calculated the proportion of studies with durations of less or equal to one generation (defined by the emergence of adults or reproduction within the experimental period), 2-5 generations, or more than five generations. We omitted 11 studies using taxa that are significantly different in life histories (e.g., insects and viruses). 109 of the remaining 257 studies (42.4%) were conducted over a time span of less than or equal to one generation and 121 studies (47.1%) lasted more than five generations of focal taxa (56, 41.2%). These results represented the slow- and fast-generating systems often used in experiments on priority effects. Indeed, studies spanning less than five generations (126, 49.0%) were conducted on mycorrhizal/wood-decomposing fungi, terrestrial/aquatic plants, amphibians, fish, and slow-generating insects (e.g., odonates), all of which require weeks to years for development and reproduction. In contrast, most studies spanning more than five generations were conducted on fast-generating microbial communities (bacteria and yeast), viruses, nematodes, trematodes, protists, phytoplankton, and zooplankton, with a small fraction of long-term studies on terrestrial plant communities (11, 4.3%). These results clearly indicate that the current body of experimental work on priority effects is strongly biased towards two extremes: short-term studies on systems with long generation times and long-term studies on systems with very short generation times (Figure IB).

In addition to system selection and experimental design, priority effects are also measured in diverse ways. The most common response variables include community composition [49], functional traits [25,26], demographic rates such as growth, survival, and fecundity [2,42,61], and ecosystem function [96,97], but studies can also measure highly specific responses such as aggression, site occupancy [28,30,98], and in coinfection experiments, pathogen load, and host response [38,99]. These measurements in part highlight the importance of priority effects in nature but also increase the difficulty of generalizing, quantifying, and comparing the consequences of priority effects across systems.

**Figure I:**
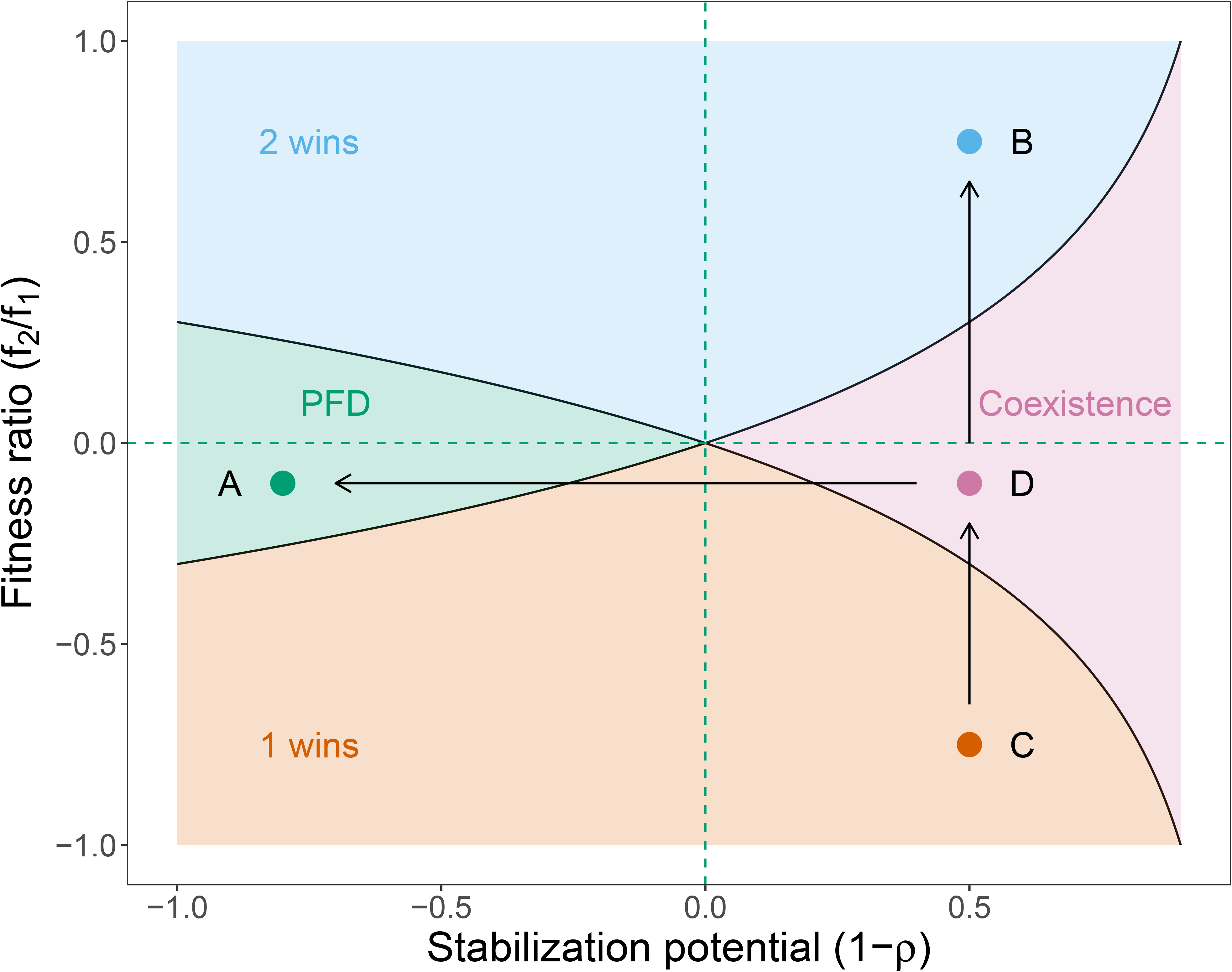
Manipulation of time in experiments and the dichotomy of time scales of experimental systems. **(A)** Results from the literature review show that 46 out of 268 experiments (17.2%) did not manipulate the arrival times of species. **(B)** Results from the literature review show the dichotomy of temporal scales among experiments: almost all studies spanning less than five generations were conducted on slow-generating systems (e.g., plants, amphibians, fish), while most studies spanning more than five generations were conducted on fast-generating systems (e.g., microbial communities, protists, planktons), with a small fraction of long-term studies on terrestrial plant communities. **(C)** Taxonomic breakdown of studies in (B). Crustaceans include zooplankton and insects.

## Box 2. Mathematical representations of trait-dependent priority effects

Trait-dependent priority effects assume a change in species interaction with their timing.

This is often modeled by a shift in interspecific competition coefficients *α*_12_ and *α*_21_, although other demographic rates may also contribute to this change. For example, consider a Beverton-Holt model where competition between two species is defined as a function of the difference between their arrival times (phenology), Δ*_p_*_12_:

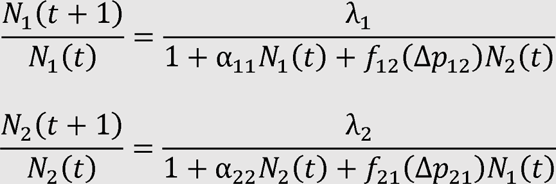

(Eqn. B2). Here, changes in the density (N_i_) between years depend on the intrinsic growth rate (*λ*) of a species which in turn is reduced by intra- and interspecific competition.

The key to the difference between frequency- and trait-dependent priority effects is termed the interaction-phenology function, α_12_ = *f*_12_(Δ*_p_*_12_), because it links per capita interaction strengths directly to their relative (typically seasonal) timing of arrival within a generation [15].

The shape of *f*(Δ*_pij_*) depends on the specific system; experiments often suggest a nonlinear relationship [2,42,46], but *f*(Δ*_pij_*) can be piecewise if species interaction only depends on whether another species is previously present, not how long it has been present (e.g., patch memory [75]; Figure 1). When the function is constant (*f*_12_(Δ*_p_*_12_) = α_12_ for all Δ*_p_*_12_), species interactions are unrelated to relative arrival times, and the model collapses to frequency-dependent priority effects (Figure 1). By accommodating a wide range of interaction-phenology relationships, this method of modeling priority effects is versatile and can be used to parameterize any empirical system.

Because both the fitness ratio and stabilization potential (*sensu* [53]) depend on interspecific competition coefficients, they also become functions of Δp_12_:

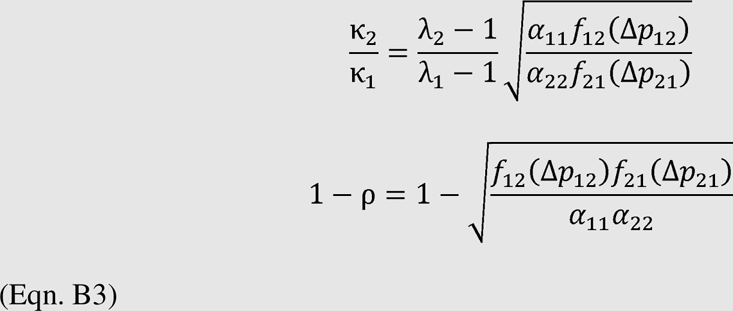

Therefore, any change in the relative timing of species would shift the position of the species pair in the coexistence space. For instance, when Δ*_p_*_12_ = 0 species 1 and 2 could coexist (point D in Figure II), but when species 1 arrives earlier (Δ*_p_*_12_ >0) the corresponding fitness ratio and stabilization potential may shift to point C, or species 1 wins. Note that these shifts may not necessarily lead to observable changes in competition outcomes, and the nature of these shifts depends on the working definition of priority effects: the arrival order changes either the outcome of species interaction (by which it would not be a trait-dependent priority effect) or the species interaction itself (by which it would). Note that *f*(Δ*_pij_*) can easily be applied to coefficients in models with other types of interactions, such as facilitation and mutualism, to calculate the corresponding fitness ratio and stabilizing potential [100,101].

**Figure I:**
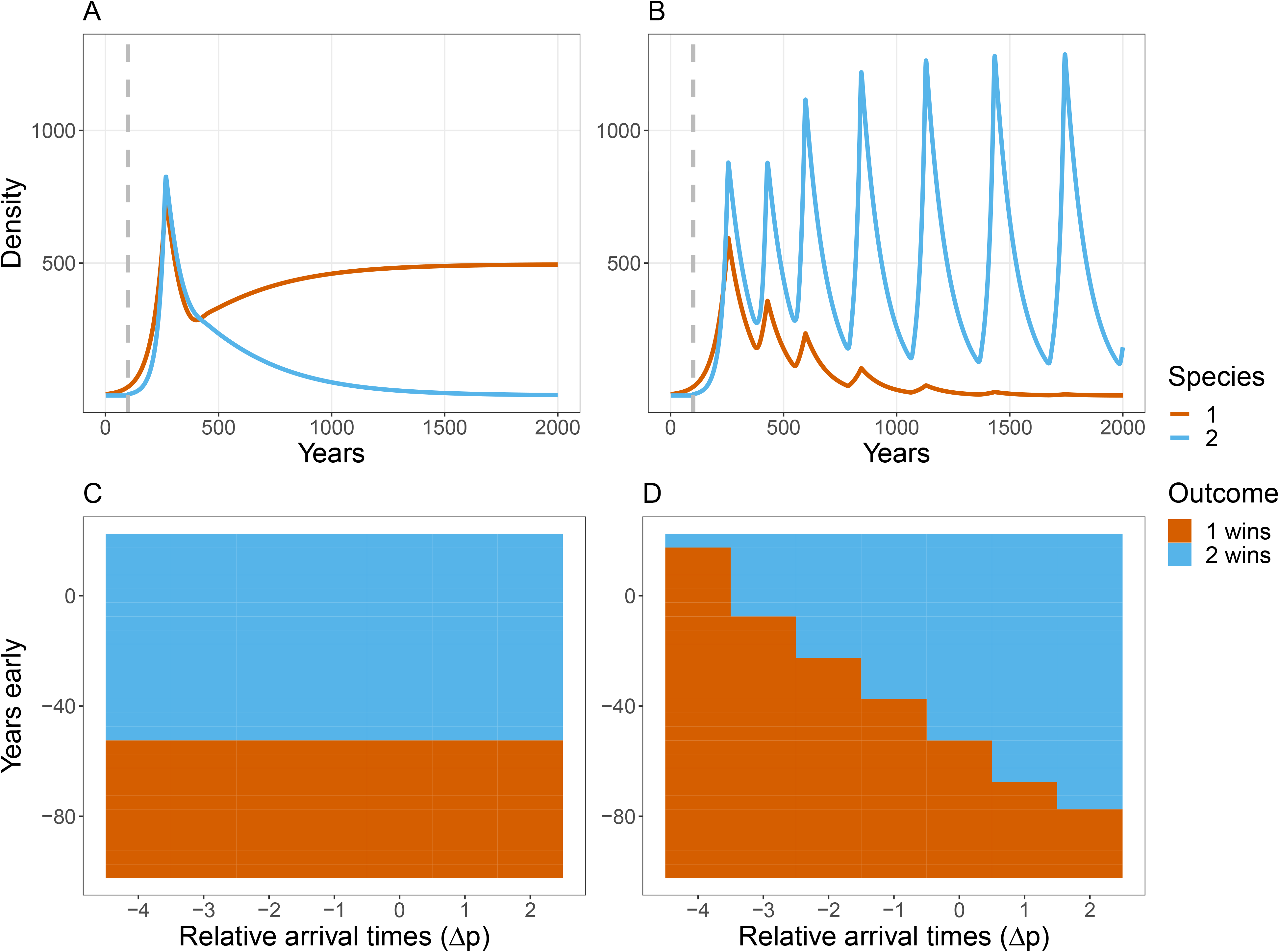
Frequency- and trait-dependent priority effects in a coexistence space. The coexistence space is often used to map species interactions with measurements of stabilizing mechanism (stabilizing potential on the x-axis) and equalizing mechanisms (fitness ratio on the y-axis). Positive frequency dependence (PFD) is represented by a region where ρ>*f*_2_/*f*_1_ >*1/ρ* (point A). Trait-dependent priority effects are defined by changes in interaction strengths due to changes in arrival times, represented by arrows. Note that a trait-dependent priority effect does not necessarily lead to exclusion; a shift from exclusion to coexistence (points C to D), or in relative abundance of species is also a trait-dependent priority effect.

## Box 3. Frequency- and trait-dependent priority effects from resource competition

Here, we illustrate the utility of our framework by unpacking a common mechanism of priority effects, resource competition [7]. In this system, the early arrival advantage could result from the higher abundance and subsequent population growth by positive frequency dependence [65]. However, the early arrival advantage could also arise from changes in the traits of individuals. For instance, the early arriver might have a larger size compared to the late arriver. This size advantage could lead to higher feeding rates in amphibians [40,102] or an advantage in light competition in plants [34].

To quantitatively capture the above examples, we analyze a modified, discrete-time version of the consumer-resource model used by Ke and Letten [53], where two consumer species compete for a shared resource. We model consumers with an annual cycle, and allow arrival times to vary within a year between species. We assume a higher feeding rate of the earlier arriver within a year. Therefore, the early arriving advantage in this model could arise from: (1) resource preemption by the early arriver, which is a numeric mechanism that leads to positive frequency dependence; (2) change in resource uptake rates, which is a trait-dependent mechanism. See Appendix III and Table S2 for model and simulation details.

When resource uptake rates are held constant, the two species display positive frequency dependence: a species wins if it arrives early over several years (Figure IA), regardless of the within-year arrival times (phenology) of the two species (Figure IC). However, when the early arriver within the year has a higher resource uptake rate, both arrival times within and across years matter: while one species could gain an advantage by establishing itself several years earlier than the other, it could still be excluded if it emerges late within a year (Figure IB, ID). These results highlight that: (1) one common biological mechanism of priority effects, resource competition, can be driven by either frequency- or trait-dependent mechanism, or a combination of both; (2) the two types of priority are not mutually exclusive, but operate on different time scales; (3) in nature, population dynamics could be affected by arrival times both within a year and across years.

**Figure I:**
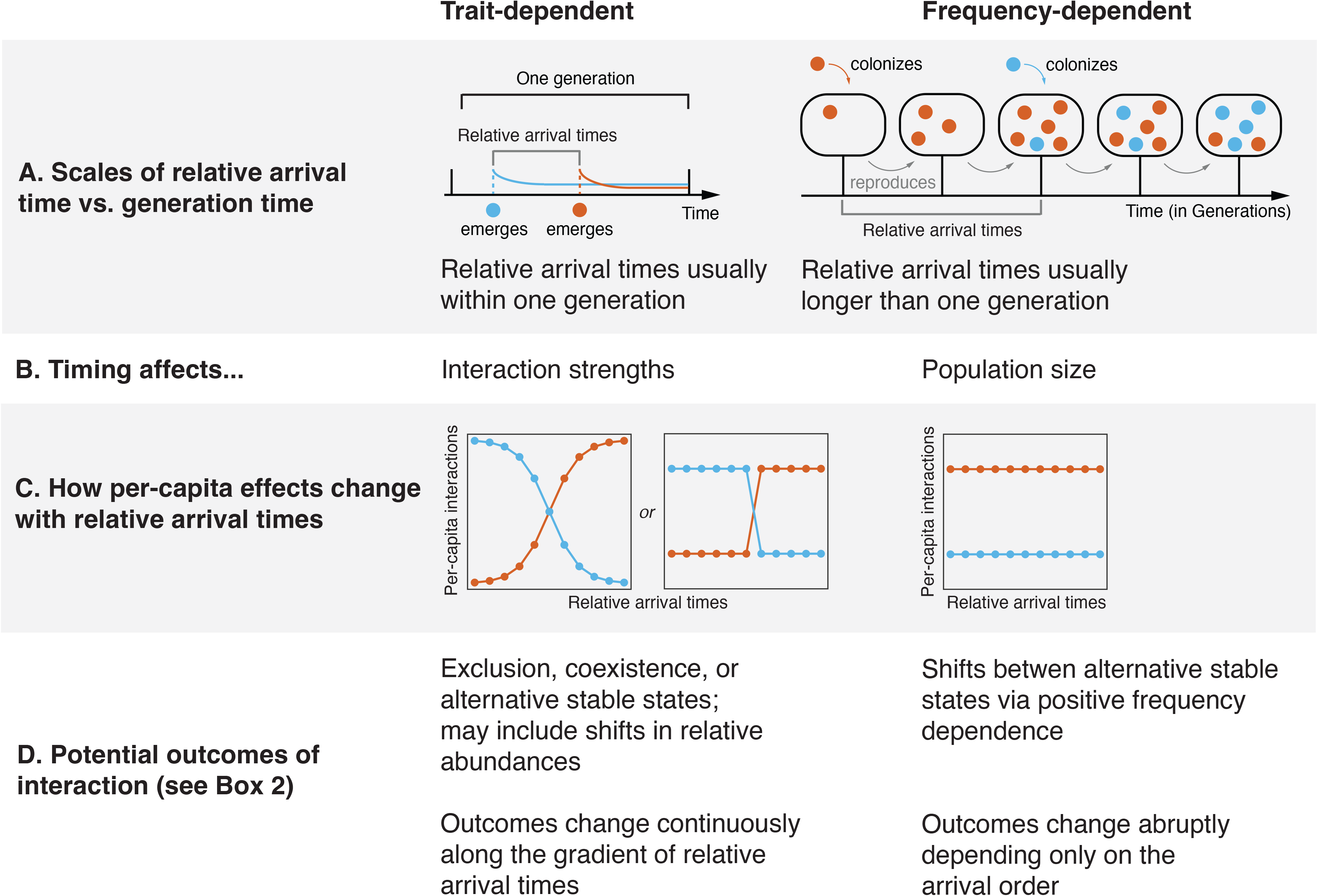
A model with both frequency- and trait-dependent priority effects. (A) Population dynamics when species 1 establishes several years earlier than species 2; species 1 has a higher density, showing a frequency-dependent priority effect. Vertical dash line indicates when species 2 arrives. (B) Population dynamics when species 1 establishes several years earlier than species 2 but emerges later than species 2 within a year; despite species 1’s frequency-dependent priority effect, species 2 has a higher feeding rate by emerging earlier within a year, displaying a trait-dependent priority effect over species 1. (C) Without trait-dependent priority effects, the relative emergence time within a year (Δ*_p_*) does not affect competition outcomes. (D) With both frequency- and trait-dependent priority effects, both the relative arrival time over years (years early on the y-axis) and within a year (Δ_*p*_, on the x-axis) affect competition outcomes.

