## Supplementary Material for "Bridging Theory and Experiments of Priority Effects"

**ORCIDs:** [1] 0000-0002-1965-6910, [2] 0000-0002-9214-2000

**Address:** [1,2] Program in Ecology and Evolutionary Biology, Department of BioSciences, Rice University, 6100 Main St, Houston, TX 77005

**List of Elements:** Table S1, Table S2; Appendix I-III

**Table S1.** Examples of experiments of priority effects, categorized by frequency-dependent or trait-dependent mechanisms, related to Figure 1.

| **Mechanism Causing or Promoting Priority Effects** | **Studies** | **System and Duration (in generations)** | **Frequency-dependent or Trait-dependent** |
|---|---|---|---|
| Size-dependent interspecific competition | [1–4] | Amphibian (1) | Trait-dependent |
| | [5] | Terrestrial plants (1) | Trait-dependent |
| Resource use overlap | [6,7] | Microbial community (>5) | ? |
| Resource depletion (before late species' arrival) | [8] | Amphibian (1) | Trait-dependent |
| Size-mediated predation/cannibalism | [9] | Amphibian (1) | Trait-dependent |
| | [10]† | Amphibian and insect larvae (1) | |
| | [11] | Odonates (1) | |
| Timing/presence of predators | [12] | Freshwater snails, tadpoles (focal) and predators (>5) | |
| | [13] | Bacterivore protists (focal) and predatory protists (>5) | |
| Absence of predator (no biotic filter) | [14] | Freshwater zooplankton (>5) | ? |
| Different response to shared pathogen | [15] | *Daphnia dentifora* (>5) | ? |
| | [16] | Terrestrial plants (1) | |
| Competition of pollinator | [17] | Terrestrial plants (1) | Trait-dependent |
| Plant-soil feedback | [5,18] | Terrestrial plants (1) | Mostly trait-dependent |
| | [19] | Terrestrial plants (1-25) | |
| Aggression towards late arriver | [20–22] | Coral reef fish (1) | Trait-dependent |
| Interference of ovipositing | [1] | Amphibian (1) | |
| High resource level in the experimental mesocosm | [18] | Terrestrial plants (1) | Likely trait-dependent |
| | [7,23] | Microbial community (>5) | ? |
| | [3] | Amphibian (1) | Trait-dependent |
| Small size of the experimental mesocosm | [24] | Freshwater protists and rotifers (>5) | Frequency-dependent |

| | | | |
|---|---|---|---|
| Low environmental variability | [25] | Microbial community (>5) | ? |
| Absence of drought | [26] | Freshwater communities (>5) | |
| High dispersal | [27,28] | Microbial community (>5) | |
| High temperature | [29] | Amphibian (1) | Trait-dependent |
| Low soil moisture | [30] | Terrestrial plants (1) | ? |
| Close phylogenetic distance between species | [6,31] | Microbial community (>5) | Likely trait-dependent |
| Early arriving exotics having larger effects on natives | [32]‡; [5] | Terrestrial plants (1) | Likely trait-dependent |
| Early flowering increases fecundity | [33] | | Trait-dependent |
| History of sympatric evolution | [34,35] | Bacteria and archaea (>5) | Trait-dependent |
| Unidentified, but calculated interspecific competition | [36] | Fungi-feeding *Drosophila* spp. (1) | Trait-dependent (calculated changes of interaction over arrival times) |
| | [37] | Freshwater protists (>5) | |
| | [38] | Terrestrial plants (1) | |
| Unidentified | [39] | Bacteria (focal) coinfecting mammal host (>5) | ? |
| | [40] | Pathogens (focal) coinfecting *Daphnia* host (>5) | |
| | [41] | Trematodes (focal) coinfecting amphibian host (1) | Trait-dependent |

‡ Did not manipulate arrival times (germination) of species.

† Manipulated initial density only.

? If original study does not provide enough evidence for categorizing mechanisms of priority effects; both are possible (e.g., the absence of predator can lead to simple numeric advantage of one prey and therefore frequency-dependent priority effects, but may also change the interactions between preys, which would lead to trait-dependent priority effects).

**Appendix I: An overview of experimental studies on priority effects, related to Box 1**

On March 24, 2022, we searched the literature on the Web of Science using the string:

*TS = (“priority effect∗”) OR TS = ("community assembly" AND ("temporal order OR “immigration history” OR “assembly order” OR “assembly history” OR “arrival order∗” OR “order of arrival∗” OR “assembly sequence∗” OR “historical contingenc∗” OR "alternative state*" OR “alternative stable state∗”)) AND SU = ("ecology" OR "evolution"),*

which returned 237 results.

Our list of articles was then compared and extended using another list of studies created by the sPriority working group funded by the Synthesis Centre of the German Research Centre for Integrative Biodiversity Research (iDiv, Germany). The sPriority group searched on the Web of Science on October 19, 2021, using an expanded list of keywords:

*(“priority effect*” OR "niche pre*empt*" OR "niche modif*" OR "historic* contingen*" OR "assembly order" OR "assembly history" OR "order of assembl*" OR “assembly sequence*” OR “sequence of assembl*" OR “sequential assembl*" OR "arrival order" OR "arrival history" OR "order of arrival*" OR “arrival sequence*” OR “sequence of arrival*" OR “sequential arrival*" OR “prior arrival*" OR "immigration order" OR "immigration history" OR "order of immigration*" OR “immigration sequence*” OR “sequence of immigration*" OR “sequential immigration*" OR “prior immigration*" OR "introduct* order" OR "introduct* history" OR "order of introduct*" OR “introduct* sequence*” OR “sequence of introduct*" OR “sequential introduc*" OR “prior introduct*" OR "infect* order" OR "infect* history" OR "order of infect*" OR “infect* sequence*” OR “sequence of infect*" OR “sequential infect*" OR “prior infect*" OR "inoculat* order" OR "inoculat* history" OR "order of inoculat*" OR “inoculat* sequence*” OR “sequence of inoculat*" OR “sequential inoculat*" OR “prior inoculat*" OR "coloni*ation* order" OR "coloni*ation* history" OR "order of coloni*ation*" OR “coloni*ation* sequence*” OR “sequence of coloni*ation*" OR “sequential coloni*ation*" OR “prior coloni*ation*" OR "invasi* order" OR "invasi* history" OR "order of invasi*" OR “invasi* sequence*” OR “sequence of invasi*" OR “sequential invasi*" OR “prior invasi*") AND*

*(experiment* OR manipulat* OR test* OR hypothes*)*

and returned 224 studies presenting results of experiments in which the order of arrival of species was manipulated. Special thanks to Chelsea Little, Judith Sarneel, Fletcher Halliday, Tamara van Steijn, Benjamin Delory and Tadashi Fukami for sharing their work with us.

Pooling both literature searches, we identified 248 studies by excluding literature from irrelevant academic fields, surveys of natural communities, mathematical models, and reviews. We also included 20 studies identified from other sources that met the search criteria but were not picked up by neither search, yielding a total of 268 studies.

For each study we identified the focal taxa (species that are manipulated), the presence and identity of secondary taxa (species that are not directly manipulated but present, such as host species in coinfection experiments), the number of species with manipulated arrival time, whether species arrive along a temporal gradient (i.e., not a simple early/late/simultaneous treatment with only three relative arrival times), duration of the experiment (in generation times of the focal taxa), main response variables measured, and presence of multiple trophic levels. See **Supplementary Data** for a list of all studies included. See https://github.com/hengxingzou/PE_Review for the code used to generate Figure I in Box 1.

**Appendix II: Mathematical representations of frequency-dependent priority effects, related to Box 2**

Consider a simple, discrete-time Beverton-Holt competition model:

$$\frac{N_1(t+1)}{N_1(t)} = \frac{\lambda_1}{1 + \alpha_{11}N_1(t) + \alpha_{12}N_2(t)}$$

$$\frac{N_2(t+1)}{N_2(t)} = \frac{\lambda_2}{1 + \alpha_{22}N_2(t) + \alpha_{21}N_1(t)}$$

(Eqn. B1).

Analytically, frequency-dependent priority effects represent a definitive outcome of pairwise competition under the modern coexistence theory framework, defined by two key parameters, fitness ratio ($\kappa_2/\kappa_1 = \frac{\lambda_2 - 1}{\lambda_1 - 1}\sqrt{\frac{\alpha_{11}\alpha_{12}}{\alpha_{22}\alpha_{21}}}$) and stabilization potential ($1 - \rho = 1 - \sqrt{\frac{\alpha_{12}\alpha_{21}}{\alpha_{11}\alpha_{22}}}$) [42,43]. Coexistence requires that the fitness ratio should be smaller than the stabilization potential, i.e., $\rho < \kappa_2/\kappa_1 < 1/\rho$ [44]. Frequency-dependent priority effects require that the reverse is true, i.e., $\rho > \kappa_2/\kappa_1 > 1/\rho$. If the stabilization potential is larger than the fitness ratio, both species cannot invade the other when rare, and the early arriver maintains its numeric advantage by positive population growth [43,45]. Therefore, this type of priority effect is particularly likely in species that have similar niches, essentially providing a mathematical formulation of the concept of "niche pre-emption" that has been proposed previously as a key mechanism of priority effects in systems dominated by competition [7,46]. With a large body of literature providing methods for quantifying competition coefficients, fitness, and niche differences [47,48] this theoretical framework allows the parameterization of competition models from experimental data to test and critically evaluate the role of priority effects on long-term structure of natural communities [16,49].

Visually, competition outcomes can be represented by a "coexistence space", where the fitness ratio is on the y-axis and stabilization potential is on the x-axis (Figure I in Box 2 of the main text). Each pairwise interaction can therefore be mapped as a point of this plane and falls into one of the three outcomes: competitive exclusion, coexistence, and positive frequency dependence (PFD).

Note that the framework of frequency-dependent priority effects does not prohibit key parameters of the competition model, such as interspecific competition coefficients, to change with relative arrival times, if the calculated fitness ratio and stabilization potential still satisfy $\rho > \kappa_2/\kappa_1 > 1/\rho$. However, the majority of models and experiments that parameterize these models assumes constant interspecific competition coefficients [23,43,45,50].

**Appendix III: Simulation details of frequency- and trait-dependent priority effects from resource competition, related to Box 3**

To quantitatively examine frequency- and trait-dependent priority effects arising from resource competition, we analyzed a modified, discrete-time version of the consumer-resource model used by Ke and Letten [43], where two consumer species ($N_1$ and $N_2$) compete for a shared resource ($R$):

$$N_1(t+1) = \lambda_1 N_1(t) \frac{R(t)^2}{k_1(\Delta p) + R(t)^2} + (1 - \mu_1) N_1$$

$$N_2(t+1) = \lambda_2 N_2(t) \frac{R(t)}{k_2(\Delta p) + R(t) + R(t)^2/k_f} + (1 - \mu_2) N_2$$

$$R(t+1) = rR(t)\left(1 - \frac{R(t)}{K}\right) + R(t) - Q_1 \lambda_1 N_1(t) \frac{R(t)^2}{k_1 + R^2} - Q_2 \lambda_2 N_2(t) \frac{1}{k_2 + R(t) + R^2(t)/k_{i2}}$$

Here, the resource grows logistically with intrinsic growth rate $r$ and carrying capacity $K$, and the resource mortality equals the consumption times a conversion factor $Q$. Consumer species 1 follows a type-III functional response, and consumer species 2 follows a type-II functional response with inhibition at higher resource levels with $k_f$ as a scaling constant determining the intensity of the inhibition. Both species are subject to density-dependent mortality ($\mu_i$).

We model consumers with an annual cycle, and allow arrival times to vary within a year between species. Within a year, each species has unique arrival time (phenology) and the phenological difference between the two species is given by $\Delta p$, which affects half-saturation constants $k_i$ of each species' functional response. We assume that the specific value of $k_i(\Delta p)$ follows the equation:

$$k_i(\Delta p) = \frac{k_{i,0}}{1 + exp\big((d - \Delta p)/c\big)}$$

Where $k_{i,0}$ is the maximum half-saturation constant and $c$ is the scaling constant that determines the shape and direction of the function. We flipped the sign of $c$ for different species to ensure that the early arriver would have a lower $k_i$, corresponding to a higher rate of resource uptake. $d$ is the value at which $k_i$ reaches half of the maximum value ($k_{i,0}$). Because the $k_1$ and $k_2$ functions are symmetric (differ by the sign of $c$), $d$ also marks where the two functions intercept, i.e., when $k_1 = k_2$. We let $d = 0$, meaning that $k_1 = k_2$ when the two species arrive simultaneously (the "midpoint"; $\Delta p = 0$). When the consumer functional response is independent of arrival times within a year (no trait-mediated mechanism), we set the value as half of the maximum ($k_{i,0}/2$); this is the calculated value of $k_i(\Delta p)$ when $\Delta p = 0$. Therefore, this model allows for differences in arrival times at two temporal scales: across years (multiple generations) and within a year; it also allows for priority effects arising from positive frequency dependence and a trait-mediated mechanism.

We simulated this model for 3,000 years, varying the arrival time over years and within each year. For each trial, we evaluated the competition outcome by a numeric method: a species is competitively excluded if it reaches a population smaller than 0.01. Preliminary trials show that this criterion is equivalent to running the model for longer but having a lower exclusion threshold. See Table 1 for all parameters used. All simulated were performed in R version 4.2.1 [51]. See https://github.com/hengxingzou/PE_Review for simulation code.

**Table S2.** Main parameter used in the simulation, related to Box 3.

| Parameter Name | Symbol | Values used |
|---|---|---|
| Intrinsic growth rate of consumer 1 | $\lambda_1$ | 0.029 |
| Intrinsic growth rate of consumer 2 | $\lambda_2$ | 0.02 |
| Mortality of the consumer | $\mu$ | 0.01 |
| Intrinsic growth rate of the resource | r | 0.5 |
| Carrying capacity of the resource | K | 3 |
| Conversion factor for both consumers 1 and 2 | $Q_1, Q_2$ | 0.01 |
| Maximum half-saturation constant of the functional response | $k_{1,0}$ | 0.02 |
| Maximum half-saturation constant of the functional response | $k_{2,0}$ | 3 |
| Inhibition constant (species 2 only) | $k_f$ | 1 |
| Half-maximum point | $d$ | 0 |
| Scaling factor for time-dependent half-saturation constant | $c_i$ | $\pm 7.5$ |
